# Agricultural landscapes with high compositional heterogeneity support both forest and farmland birds in Taiwan

**DOI:** 10.1101/2024.07.16.603643

**Authors:** Da-Li Lin, Tatsuya Amano, Richard A. Fuller, Tzung-Su Ding, Martine Maron

## Abstract

**Context:** Promoting heterogeneous agricultural landscapes could help to reduce the negative impacts of habitat conversion on biota. However, the benefits of landscape heterogeneity can vary among spatial scales and taxa.

**Objectives:** To design biodiversity-friendly landscapes, we use nationwide bird survey data and land use maps to examine the effects of compositional heterogeneity, configurational heterogeneity, and habitat amount at different scales on the species richness of different bird groups.

**Methods:** We examine the effects of configurational heterogeneity (measured using edge density), compositional heterogeneity (Shannon’s diversity index of habitat types), and habitat amount (proportion of forest and farmland cover) at both transect (local) and landscape (0.5, 1, or 2 km) scales on the species richness of all breeding birds, forest birds, farmland birds, and introduced birds.

**Results:** Total species richness had a hump-shaped relationship with local forest cover, and with farmland cover at landscape scale. Richness of both forest birds and richness of farmland birds increased with Shannon’s diversity index of habitat types at both local and landscape scales, but only increased with the amount of their preferred habitat at the local scale. Richness of introduced birds was greater in landscapes with higher edge density, suggesting those species are associated with human-dominated landscapes.

**Conclusions:** High compositional heterogeneity with low configurational heterogeneity at the landscape scale may help maintain native bird richness while minimising the spread of introduced species in Taiwan. These results can help guide land use planning to achieving biodiversity goals in a country with intensive land-use competition.

## 1. Introduction

Agricultural expansion and intensification are among the primary drivers of biodiversity loss (Tilman et al. 2001; Foley et al. 2011; Maxwell et al. 2016). Many species that rely on natural habitats are negatively affected by conversion to built-up areas and/or intensively-managed agricultural lands (e.g., Stanton et al. 2018). For example, in recent decades, agricultural practices have been causing serious farmland bird declines in North America (Kirk et al. 2011) and Europe (Newton 2004). Declining farmland biodiversity may further affect ecosystem functions and the provision of ecosystem services, including crop production, pest control, and pollination (Tscharntke et al. 2005; Zhang et al. 2007).

Under some circumstances, agricultural landscapes can support rich biodiversity and thus the provision of a wide range of ecosystem functions and services (Power 2017). Agricultural lands can be highly heterogeneous, with numerous types of habitats (Benton et al. 2003; Fahrig et al. 2011; Kremen & Merenlender 2018). Agricultural landscapes that contain a complex mosaic of diverse land cover types usually support higher biodiversity than those with a few types, or monocultures (Fischer & Lindenmayer 2007; Mendenhall et al. 2014). Increasing the heterogeneity of agricultural landscapes can help conserve biodiversity on farmlands (Kremen & Merenlender 2018).

Identifying the associations between landscape heterogeneity and diversity of target taxa in agricultural landscapes can help design management approaches (Andrén 1994; Benton et al. 2003; Tscharntke et al. 2005). Based on the concept of functional landscape heterogeneity, compositional heterogeneity and configurational heterogeneity are two important metrics of landscapes (Fahrig & Nuttle 2005; Fahrig et al. 2011). Compositional heterogeneity refers to the diversity of land cover types, while configurational heterogeneity reflects the spatial complexity of a given landscape (Fahrig & Nuttle 2005). Changes in either or both metrics can influence local biodiversity, and the associations usually vary geographically and among taxa (e.g., Perović et al. 2015; Neumann et al. 2016; Yang et al. 2021).

Although the loss of natural habitats due to conversion to agriculture has generally negative impacts on local biodiversity (e.g., Guerrero-Pineda et al. 2022), even small patches of retained habitat can help maintain some wildlife populations and provide ecosystem services (e.g., Tulloch et al. 2016; Riva & Fahrig 2022). Indeed, Fahrig (2013) suggested that the total amount of habitat in a given landscape unit is more important than its configuration in predicting species richness. Yet, because the association between habitat amount and species richness is not consistent (e.g., not supported by Haddad et al. 2017; Saura 2021 while supported by Merckx et al. 2019), local investigations are needed to confirm the situation in any particular location and context.

Moreover, among spatial scales, the relationship between landscape heterogeneity and biodiversity can change or even reverse (DeFries et al. 2010; Morelli et al. 2013; Schindler et al. 2013), indicating that the effectiveness of landscape management actions may depend crucially on spatial scale. That is, by conducting management actions at inappropriate scale, the results for biodiversity conservation could weaken (e.g., Pickeet and Siriwadena 2011). Agri-environmental schemes thus need to work at the most appropriate scale for target taxa, or even design actions at multiple scales to maximize the effectiveness of biodiversity conservation (Gabriel et al. 2010).

Because Taiwan is a mountainous island with limited flat land there is high land use competition among different sectors (Chung 2019). In recent years, Taiwan’s authorities and stakeholders in the agricultural sector have expressed interest in integrating sustainable agricultural management with biodiversity conservation (e.g., Chang 2013; Lee et al. 2016). To design biodiversity-friendly landscapes, it is necessary to clarify how different elements of landscape heterogeneity at different spatial scales affect local bird richness. Here we use nationwide bird survey data and land use maps to examine the effects of compositional heterogeneity, configurational heterogeneity, and habitat amount at different scales on the species richness of different bird groups detected on transects. These results can inform landscape planning for biodiverse agricultural landscapes in Taiwan.

## 2. Methods

### 2.1 Study area

The study area was Taiwan island (its outlying islands were not included), a mountainous sub-tropical island (ca. 36,104 km^2^) in East Asia. The western one-third is dominated by flat lands and foothills, and the eastern two-thirds is dominated by five forest-covered mountain ranges with about 280 peaks >3,000 m above sea level. Extreme topographic and climate heterogeneity leads to several distinct ecosystem types in a relatively small area. The lowland and submontane zones (ca. 500-1,800 m) are covered by subtropical evergreen broad-leaved forests, the main montane zone (ca. 1,800-2,400 m) is covered by deciduous and evergreen broad-leaved forests, and the upper-montane zone (ca. 2,400-3,600m) is home to coniferous forests (Li et al. 2013). A mosaic of micro-habitats exists amongst these ecosystems, supporting an array of endemic species. Overall, the landscape of Taiwan is dominated by forest (58%) and farmland (20%, including rice fields (4%), orchards (5%), vegetable and coarse cereals (4%), and others (7%)), with smaller areas of coverage by water bodies (1.6%) and built infrastructure (12%; Ministry of the Interior of Taiwan 2015). With a high human population density and limited availability of flat land, competition for land-use is intensive among various sectors in Taiwan (Chung 2019).

### 2.2 Bird survey data

The Taiwan Breeding Bird Survey (BBS Taiwan) is a nationwide citizen science project that aims to monitor breeding bird population status and trends to inform conservation management (Fig. 1a, Ko et al. 2017). It was launched in 2011 as a partnership between the Endemic Species Research Institute, the Institute of Ecology and Evolutionary Biology of National Taiwan University and the Taiwan Wild Bird Federation. The BBS Taiwan comprises transects of 1-2 km in length, located according to a random stratified design (Ko et al. 2017). Taiwan Island was divided into 91 regions, based on 41 ecoregions (Su 1992) and three elevation ranges (0–1,000, 1,001–2,500, 2,501–4,000 m). Grid cells (1 km × 1 km squares) without roads or trails in good condition were excluded to ensure ease of accessibility by volunteers, and 428 cells were randomly selected from the remainder, with the number of selected grid cells being proportional to the area of the region in which they occur (Ko et al. 2017). A transect was laid out in each selected grid cell by the surveyor who first adopted it.

**Fig. 1.**
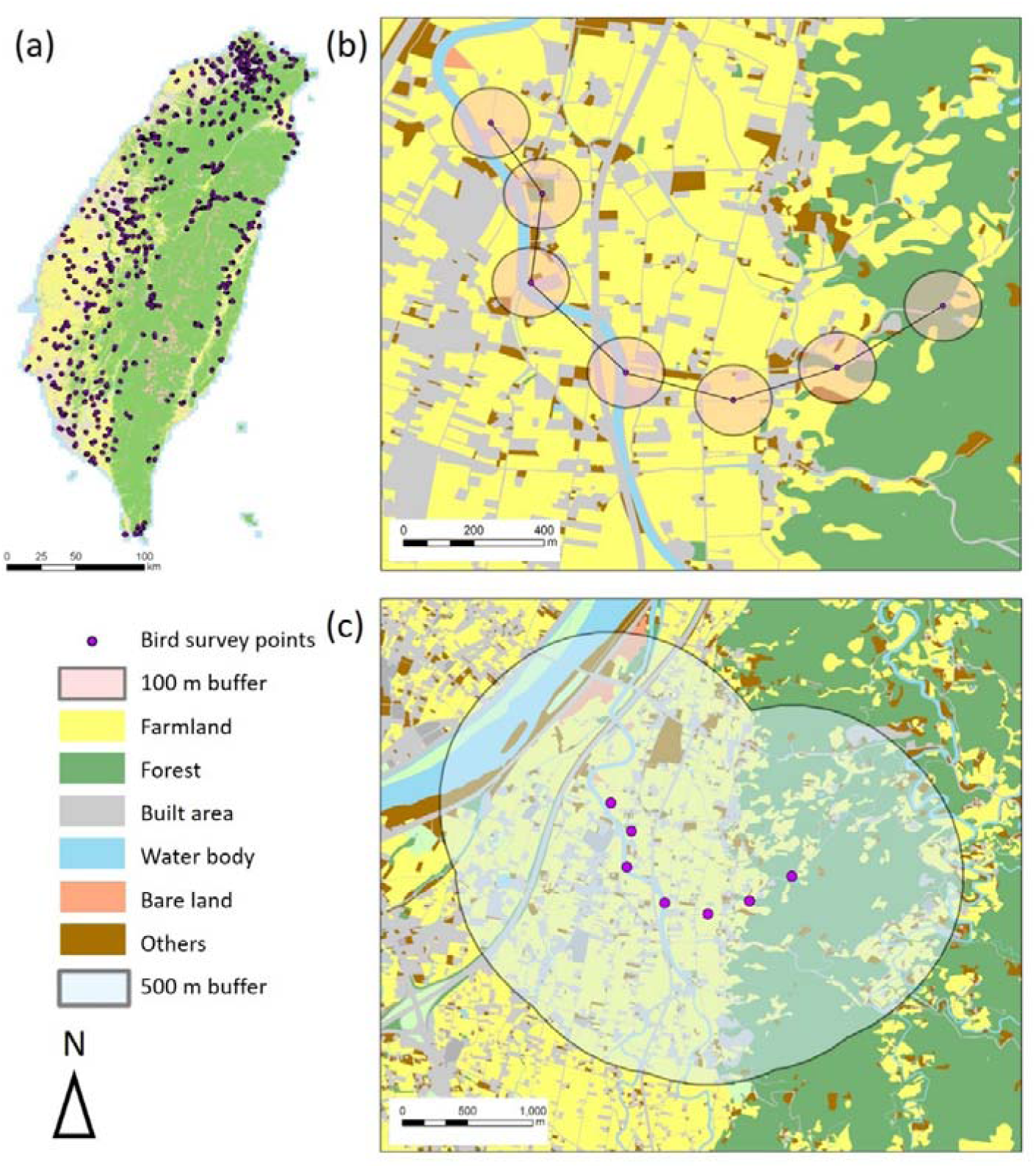
(a) Map of the main land use types and the bird transects of Taiwan Breeding Bird Survey; (b) example of the spatial arrangement of survey areas within a transect (shown with a black line); (c) an example of a transect buffer (0.5-km) around a set of survey points within which landscape context variables were calculated.

Each transect contains six to ten survey points spaced at least 200 m apart (see an example in Fig. 1b, the transect (black line) comprising seven survey points (purple)). A trained surveyor visited the points along each transect twice each year within 4 hours after sunrise on days with mild weather during the breeding season (March to June inclusive). The two visits were at least four weeks apart. On each visit to a survey point, the surveyor conducted a 6-minute stationary point count and recorded all the birds detected by sight or sound within a 100 m radius of point (the survey area). For each transect, the richness of (i) total breeding bird species, (ii) forest birds, (iii) farmland birds, and (iv) introduced birds were calculated across all survey areas that comprise the transect, and these transect-level measures were used as the response variable (Lin et al. 2023).

### 2.3 Land use data

We characterised the landscape using a high spatial resolution land use map (at the metre scale; Ministry of the Interior of Taiwan 2015; Fig. 1), reclassifying the mapped land use types into six broad classes, including farmlands (any land use patches that are managed for growing crops, excluding orchards), forests (all native and artificial woodland patches), built-up areas, water bodies, bare lands, and others. To quantify habitat amount, we measured the land use within each 100-m radius survey area within a transect (translucent circular areas in Fig. 1b) and took the average proportion of forest (as local habitat amount for forest birds) and the average proportion of farmland (as local habitat amount for farmland birds) within survey areas for each transect. We also measured the proportion of forest (as habitat amount for forest birds at landscape level) and the proportion of farmland (as habitat amount for farmland birds at landscape level) within 0.5-, 1-, and 2-km buffers around each transect (0.5 km radius example shown as pale translucent areas in Fig. 5-1c). In the transect buffers (0.5-km radius), mean elevation (“Elev”) and the area of buffer (“Area”) were also calculated. Shannon’s diversity index (H’) of land cover types 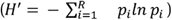 was used to index compositional heterogeneity within each of buffers, where *pi* is the proportion of land cover type *i*. Edge density (“Edge”) was used to index configurational heterogeneity; to do this, we divided the total perimeter length of each land use patch (excluding built-up area) by the area of each transect buffer. All explanatory variables are listed in Table 1.

**Table 1.** Explanatory variables in generalised linear mixed models.

| Variable code | Definition | Scale at which measured | Represents |
| --- | --- | --- | --- |
| Forest<sub>p</sub> | Mean proportion of forest cover across all sites within each transect | Survey area (local) | Habitat amount for forest birds |
| Farm<sub>p</sub> | Mean proportion of farmland cover across all sites within each transect | Survey area (local) | Habitat amount for farmland birds |
| Elev | Mean elevation | Landscape (0.5-, 1-, 2-km buffer around each transect) |  |
| Area | Area of buffer | Landscape (0.5-, 1-, 2-km buffer around each transect) |  |
| H' | Shannon diversity index of land uses | Landscape (0.5-, 1-, 2-km buffer around each transect) | Configurational heterogeneity |
| Edge | Edge density | Landscape (0.5-, 1-, 2-km buffer around each transect) | Compositional heterogeneity |
| Forest<sub>T</sub> | Proportion of landscape-scale forest cover | Landscape (0.5-, 1-, 2-km buffer around each transect) | Habitat amount for forest birds |
| Farm<sub>T</sub> | Proportion of landscape-scale farmland cover | Landscape (0.5-, 1-, 2-km buffer around each transect) | Habitat amount for farmland birds |

### 2.4 Statistical analysis

We used generalised linear mixed models with a Poisson distribution using the R package *lme4* (Bates et al. 2015) in R version 4.0.2 (R Core Team 2020) to examine how transect-level bird species richness was related to the eight explanatory variables shown in Table 1. The richness of all bird species, forest bird species, farmland bird species, and invasive bird species were the response variables. For the two local-scale explanatory variables (Forest_P_ & Farm_P_) we included both linear and quadratic terms. For the two landscape-scale explanatory variables (Forest_T_ & Farm_T_), we included both linear and quadratic terms, and tested for associations separately at the 0.5-, 1-, and 2-km scales. For Shannon’s diversity index and edge density we included linear term only (Burnham & Anderson 2002), and tested for associations separately at the 0.5-, 1-, and 2-km scales. All explanatory variables were standardised to z-scores prior to analysis.

To determine the spatial scale at which each of the four landscape-scale explanatory variables best explained variation in species richness, we developed six alternative models for each response variable. Each included Elev and Area, but different models included proportion of forest and proportion of forest farmland calculated within the 0.5-, 1-, or 2-km buffer around each transect, only as a linear term, or as both linear and quadratic terms. We then used the model with the lowest Akaike’s Information Criterion (AIC) to determine the best spatial scale and form for the explanatory variable (e.g., linear and quadratic terms at 0.5-km scale in Table 2).

**Table 2.** Statistical analysis summary of generalised linear mixed models of each species group. SE: standard error.

|  | Term | Scale | coefficient | SE | Z value | P value |
| --- | --- | --- | --- | --- | --- | --- |
| ***Total bird species richness*** |  |  |  |  |  |  |
| Forest<sub>p</sub> | L & Q | local | -0.076 | 0.018 | -4.27 | < 0.001 |
| Farm<sub>p</sub> | L | local | 0.059 | 0.016 | 3.08 | < 0.001 |
| Elev | L | 0.5 km | 0.039 | 0.012 | 3.30 | < 0.001 |
| Area | L | 0.5 km | 0.066 | 0.010 | 6.98 | < 0.001 |
| Edge | L | 1 km | -0.082 | 0.013 | -6.59 | < 0.001 |
| H' | L | 1 km | 0.052 | 0.016 | 3.17 | < 0.01 |
| Forest<sub>T</sub> | L & Q | 0.5 km | -0.036 | 0.026 | -1.42 | 0.155 |
| Farm<sub>T</sub> | L & Q | 2 km | -0.041 | 0.011 | -3.90 | < 0.01 |
| ***Forest bird species richness*** |  |  |  |  |  |  |
| Forest<sub>p</sub> | L | local | 0.091 | 0.032 | 2.88 | < 0.01 |
| Farm<sub>p</sub> | L | local | -0.196 | 0.041 | -4.83 | < 0.001 |
| Elev | L | 0.5 km | 0.106 | 0.018 | 6.04 | < 0.001 |
| Area | L | 0.5 km | 0.035 | 0.016 | 2.26 | < 0.05 |
| Edge | L | 2 km | 0.028 | 0.034 | 0.84 | 0.402 |
| H' | L | 1 km | 0.216 | 0.023 | 9.19 | < 0.001 |
| Forest<sub>T</sub> | L | 2 km | 0.463 | 0.046 | 10.03 | < 0.001 |
| Farm<sub>T</sub> | L & Q | 1 km | -0.072 | 0.025 | -2.86 | < 0.01 |
| ***Farmland bird species richness*** |  |  |  |  |  |  |
| Forest<sub>p</sub> | L | local | -0.394 | 0.040 | -9.74 | < 0.001 |
| Farm<sub>p</sub> | L | local | 0.144 | 0.027 | 5.29 | < 0.001 |
| Elev | L | 0.5 km | -0.157 | 0.032 | -4.85 | < 0.001 |
| Area | L | 0.5 km | 0.102 | 0.019 | 5.27 | < 0.001 |
| Edge | L | 1 km | -0.108 | 0.028 | -3.81 | < 0.001 |
| H' | L | 1 km | 0.152 | 0.032 | 4.78 | < 0.001 |
| Forest<sub>T</sub> | L & Q | 0.5 km | -0.138 | 0.047 | -2.91 | < 0.01 |
| Farm<sub>T</sub> | L & Q | 2 km | -0.021 | 0.018 | -1.14 | 0.253 |
| ***Introduced bird species richness*** |  |  |  |  |  |  |
| Forest<sub>p</sub> | L | local | -0.503 | 0.081 | -6.19 | < 0.001 |
| Farm<sub>p</sub> | L | local | 0.083 | 0.054 | 1.54 | 0.122 |
| Elev | L | 0.5 km | -0.563 | 0.088 | -6.44 | < 0.001 |
| Area | L | 0.5 km | 0.072 | 0.038 | 1.88 | 0.060 |
| Edge | L | 2 km | 0.180 | 0.059 | 3.05 | < 0.01 |
| H' | L | 2 km | -0.080 | 0.054 | -1.49 | 0.136 |
| Forest<sub>T</sub> | L & Q | 2 km | -0.175 | 0.065 | -2.69 | < 0.01 |
| Farm<sub>T</sub> | L | 2 km | -0.213 | 0.051 | -4.16 | < 0.001 |

Abbreviations: L: linear term, Q: quadratic term, Edge: edge density, H': Shannon's diversity index, Elev: mean elevation, Area: area of buffer, BSR: bird species richness, NS: non-significant

After identifying the best spatial scale and form for each of the six explanatory variables for each response variable, we developed the full model with all six explanatory variables calculated at the best spatial scale and in the best form, two other explanatory variables (Elev and Area) using a GLMM with a Poisson distribution with the package *lme4* (Bates et al. 2015) in R version 4.0.2 (R Core Team 2020). To assess multicollinearity among explanatory variables in the full model, we calculated Variance Inflation Factors (VIF) scores using the package *car* (Fox & Weisberg 2019). If all VIF scores were smaller than five, indicating a relatively weak effect of collinearities among the explanatory variables.

## 3. Results

The comparison of models with each explanatory variable calculated at different spatial scales and in different forms is shown in Tables S2, S3, S4, and S5. All VIF scores were smaller than five, indicating relatively weak collinearity among explanatory variables (Table S6). The correlation matrices of each species group are shown in S7, S8, S9, and S10. The analyses summaries of GLMM of each species group are listed in Table 3.

**Table 3.**
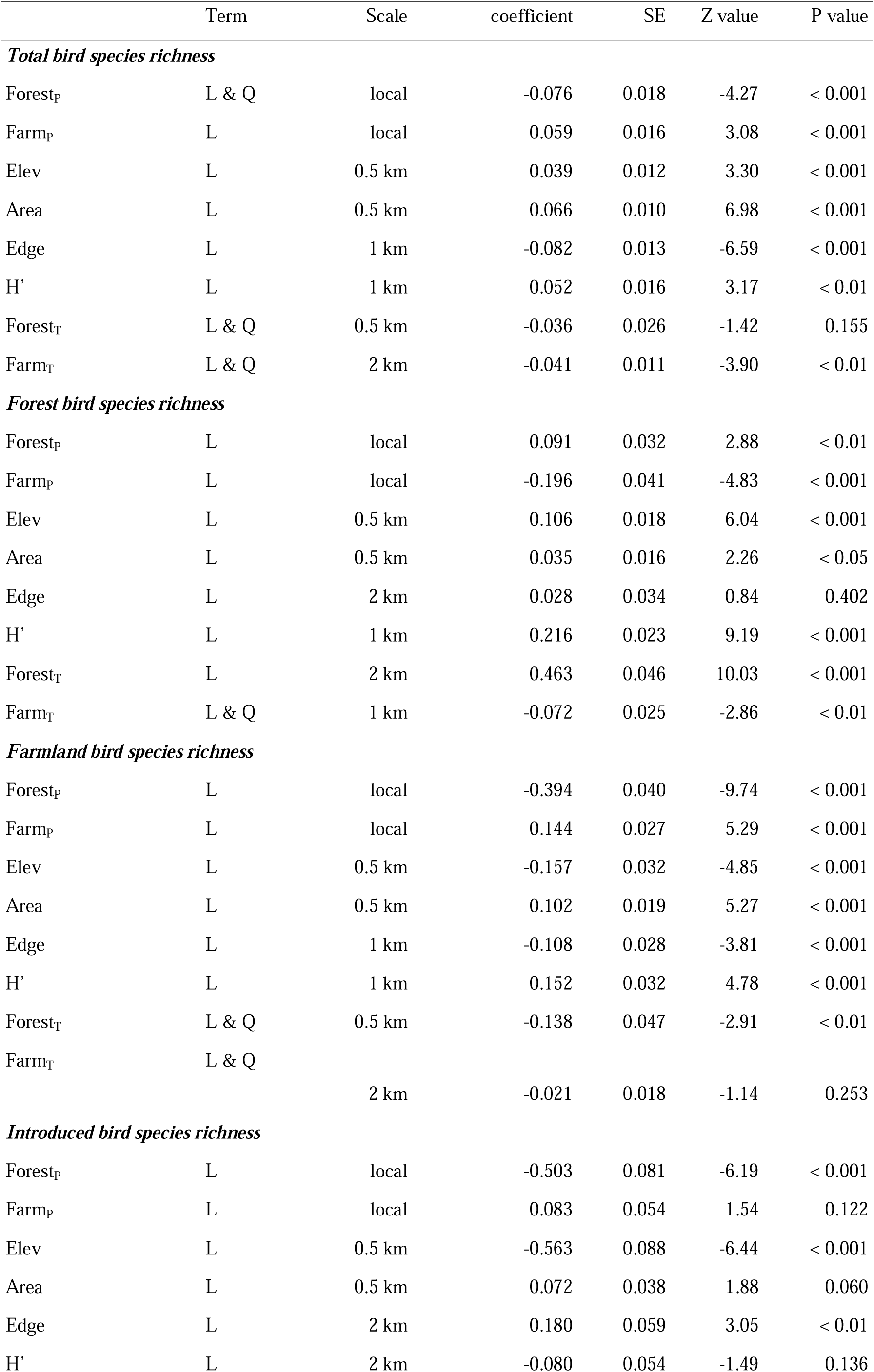
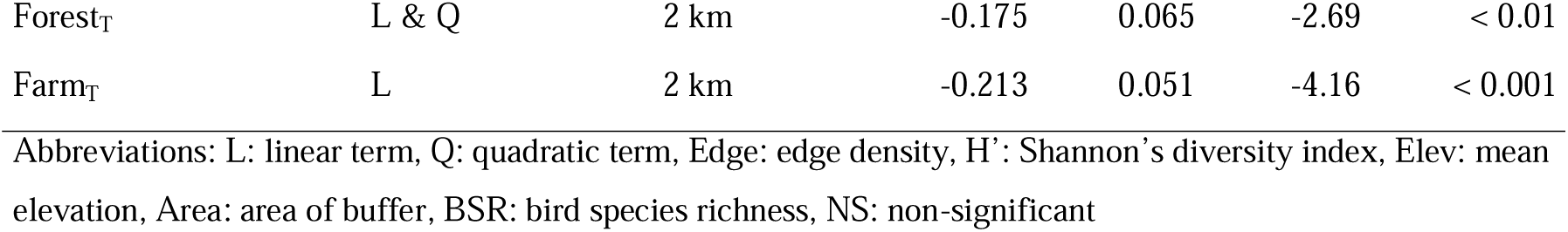
Statistical analysis summary of generalised linear mixed models of each species group. SE: standard error.

|  | Term | Scale | coefficient | SE | Z value | P value |
| --- | --- | --- | --- | --- | --- | --- |
| ***Total bird species richness*** |  |  |  |  |  |  |
| Forest<sub>p</sub> | L & Q | local | -0.076 | 0.018 | -4.27 | < 0.001 |
| Farm<sub>p</sub> | L | local | 0.059 | 0.016 | 3.08 | < 0.001 |
| Elev | L | 0.5 km | 0.039 | 0.012 | 3.30 | < 0.001 |
| Area | L | 0.5 km | 0.066 | 0.010 | 6.98 | < 0.001 |
| Edge | L | 1 km | -0.082 | 0.013 | -6.59 | < 0.001 |
| H' | L | 1 km | 0.052 | 0.016 | 3.17 | < 0.01 |
| Forest<sub>T</sub> | L & Q | 0.5 km | -0.036 | 0.026 | -1.42 | 0.155 |
| Farm<sub>T</sub> | L & Q | 2 km | -0.041 | 0.011 | -3.90 | < 0.01 |
| ***Forest bird species richness*** |  |  |  |  |  |  |
| Forest<sub>p</sub> | L | local | 0.091 | 0.032 | 2.88 | < 0.01 |
| Farm<sub>p</sub> | L | local | -0.196 | 0.041 | -4.83 | < 0.001 |
| Elev | L | 0.5 km | 0.106 | 0.018 | 6.04 | < 0.001 |
| Area | L | 0.5 km | 0.035 | 0.016 | 2.26 | < 0.05 |
| Edge | L | 2 km | 0.028 | 0.034 | 0.84 | 0.402 |
| H' | L | 1 km | 0.216 | 0.023 | 9.19 | < 0.001 |
| Forest<sub>T</sub> | L | 2 km | 0.463 | 0.046 | 10.03 | < 0.001 |
| Farm<sub>T</sub> | L & Q | 1 km | -0.072 | 0.025 | -2.86 | < 0.01 |
| ***Farmland bird species richness*** |  |  |  |  |  |  |
| Forest<sub>p</sub> | L | local | -0.394 | 0.040 | -9.74 | < 0.001 |
| Farm<sub>p</sub> | L | local | 0.144 | 0.027 | 5.29 | < 0.001 |
| Elev | L | 0.5 km | -0.157 | 0.032 | -4.85 | < 0.001 |
| Area | L | 0.5 km | 0.102 | 0.019 | 5.27 | < 0.001 |
| Edge | L | 1 km | -0.108 | 0.028 | -3.81 | < 0.001 |
| H' | L | 1 km | 0.152 | 0.032 | 4.78 | < 0.001 |
| Forest<sub>T</sub> | L & Q | 0.5 km | -0.138 | 0.047 | -2.91 | < 0.01 |
| Farm<sub>T</sub> | L & Q |  |  |  |  |  |
|  |  | 2 km | -0.021 | 0.018 | -1.14 | 0.253 |
| ***Introduced bird species richness*** |  |  |  |  |  |  |
| Forest<sub>p</sub> | L | local | -0.503 | 0.081 | -6.19 | < 0.001 |
| Farm<sub>p</sub> | L | local | 0.083 | 0.054 | 1.54 | 0.122 |
| Elev | L | 0.5 km | -0.563 | 0.088 | -6.44 | < 0.001 |
| Area | L | 0.5 km | 0.072 | 0.038 | 1.88 | 0.060 |
| Edge | L | 2 km | 0.180 | 0.059 | 3.05 | < 0.01 |
| H' | L | 2 km | -0.080 | 0.054 | -1.49 | 0.136 |
| Forest<sub>T</sub> | L & Q | 2 km | -0.175 | 0.065 | -2.69 | < 0.01 |
| Farm<sub>T</sub> | L | 2 km | -0.213 | 0.051 | -4.16 | < 0.001 |

The full model showed a significant hump-shaped association between local forest cover and species richness of all breeding birds (Fig. 2a1, p < 0.001), though forest cover was not significant at the landscape (0.5-km) scale (Fig. 2a5, p = 0.155). On the other hand, species richness of all breeding birds was significantly negatively associated with farmland cover at the local scale (Fig. 2a2, p < 0.001), and showed a significant hump-shaped association at the landscape (2-km) scale (Fig. 2a6, p < 0.001). Species richness of all breeding birds also showed a significant negative association with edge density (Fig. 2a3, p < 0.001) and a significant positive association with Shannon’s diversity index (Fig. 2a4, p < 0.01). Both mean elevation (Fig 2a7, p < 0.001) and area of buffer (Fig. 2a8, p < 0.001) were significantly positively associated with species richness of all breeding birds.

**Fig 2.**
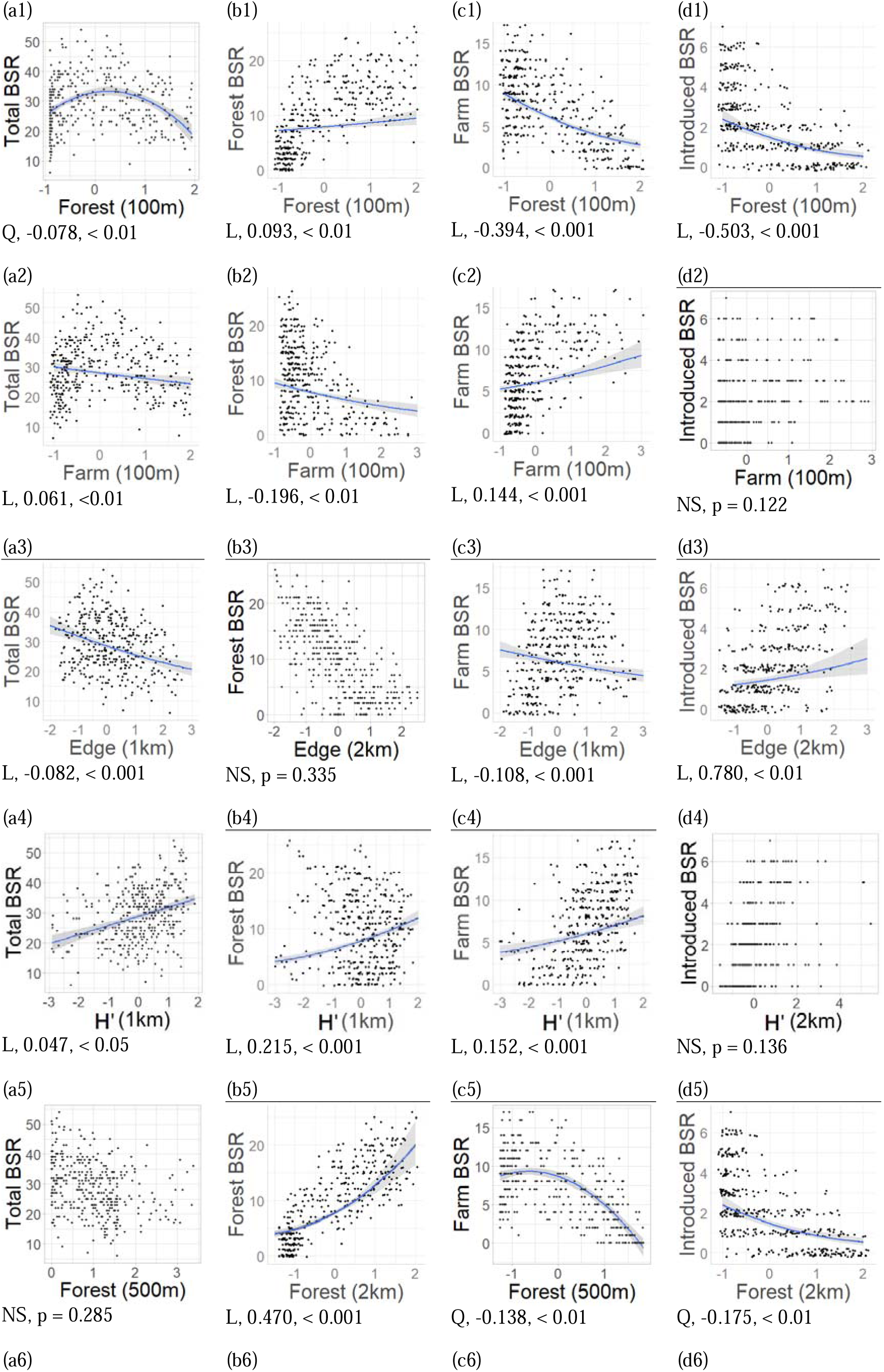
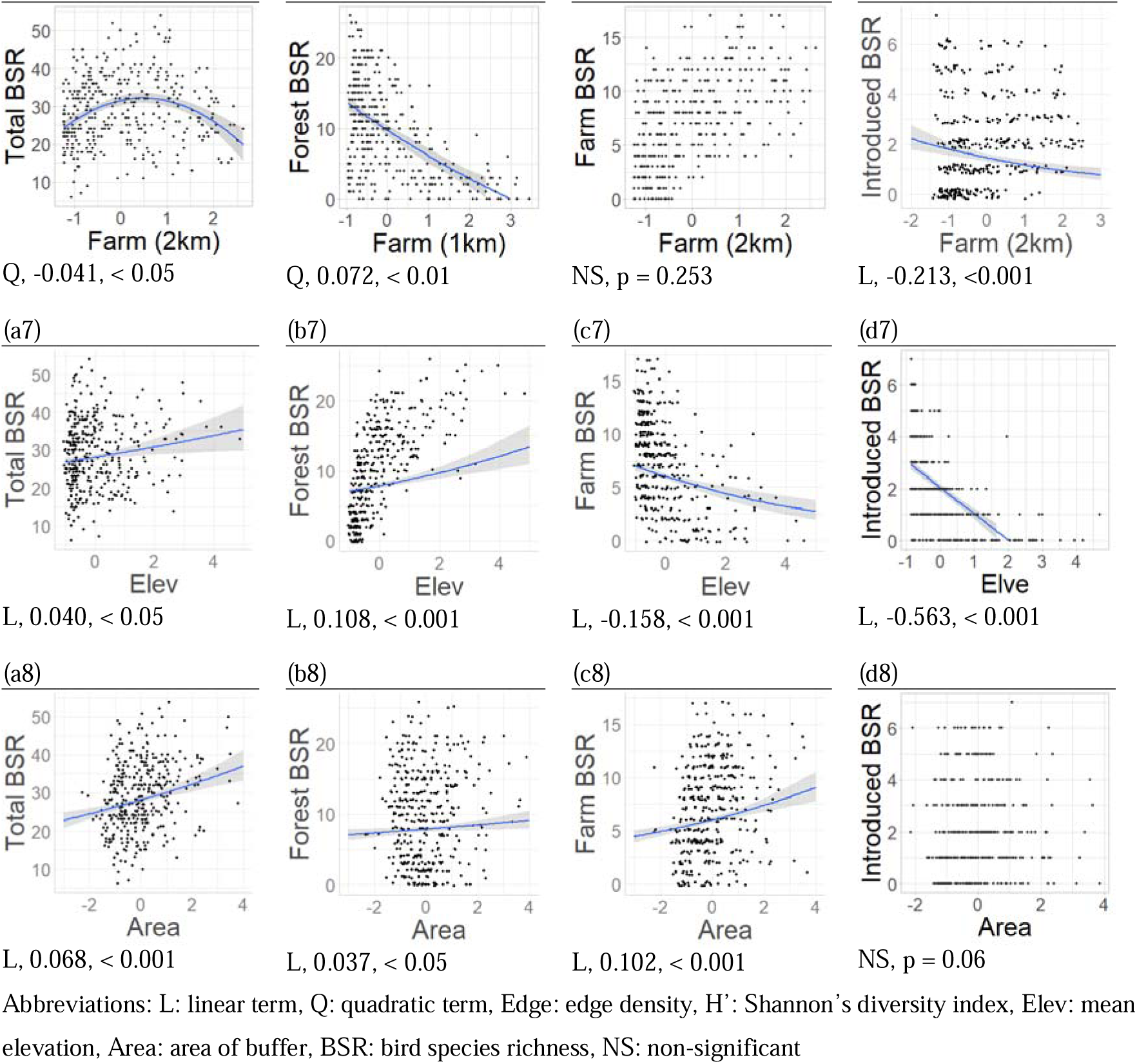
Relationships between bird species richness and explanatory variables. Regression lines are shown only for significant variables, based on the coefficients estimated with generalised linear mixed models using both linear and quadratic terms. The numbers below the figure indicate coefficients and p-values.

Forest cover was significantly positively associated with forest bird species richness at both local (Fig. 2b1, p < 0.01) and landscape (2-km, Fig. 2b5, p < 0.001) scales. In contrast, farmland cover was significantly negatively associated with forest bird species richness at both local (Fig. 2b2, p < 0.001) and landscape (1-km, Fig. 2b6, in quadratic term, p < 0.01) scales. Edge density was not significantly associated with forest bird species richness (Fig. 2b3, p = 0.402), but Shannon’s diversity index was significantly positively associated with forest bird species richness (Fig. 2b4, p < 0.001). Both mean elevation (Fig. 2b7, p < 0.001) and area of buffer (Fig. 2b8, p < 0.05) were significantly positively associated with forest bird species richness.

For farmland bird species richness, forest cover at both local (Fig. 2c1, p < 0.001) and landscape scales (Fig. 2c5, in quadratic term, p < 0.001) was significantly negatively associated. In contrast, the association with farmland cover was significantly positive at the local scale (Fig. 2c2, p < 0.001) but not significant at the landscape (2-km) scale (Fig. 2c6, p = 0.253). Farmland bird species richness was also significantly negatively associated with edge density (Fig. 2c3, p < 0.001) and positively with Shannon’s diversity index (Fig. 2c4, p < 0.001). Moreover, mean elevation was significantly negatively associated with species richness of farmland birds (Fig. 2c7, p < 0.001) and area of the buffer was significantly positively associated (Fig. 2c8, p < 0.001).

For introduced bird species richness, forest cover at local (Fig. 2d1, p < 0.001) and landscape (2-km) scales (Fig. 2d5, quadratic term, p < 0.01) showed significantly negative associations. On the other hand, farmland cover was not significantly associated at the local scale (Fig. 2d2, p = 0.122) but was negatively associated at the landscape (2-km) scale (Fig 2d6, p < 0.001). In contrast to other species groups, the association between richness of introduced birds and edge density was significantly positive (Fig. 2d3, p < 0.01), and there was no significant association between richness and Shannon’s diversity index (Fig. 2d4, p = 0.136). Mean elevation was significantly negatively associated with introduced bird species richness (Fig. 2d7, p < 0.001) while area of buffer had no significant association (Fig. 2d8, p = 0.06).

## 4. Discussion

### 4.1 Compositional heterogeneity supports high bird diversity

The results of this study suggest that a landscape with high compositional heterogeneity and low configurational heterogeneity may support high breeding bird species richness in Taiwan, especially where both forest and farmland patches are present. For richness of all breeding birds, the hump-shaped associations of local forest cover (Fig. 2a1) and landscape farmland cover (Fig. 2a6) suggest that an intermediate proportion of forest and farmland cover (agriculture-forest mosaic), rather than landscapes dominated by forest and farmland, supports most breeding bird species. These results are similar to the case in Japan (Katayama et al. 2014) and support the concept of functional landscape heterogeneity (Fahrig et al. 2011) and working lands conservation (Kremen & Merenlender 2018), in which a mosaic landscape comprising forest and farmland patches supports more species than a single large forest or farmland patch. Moreover, generalist species can also presumably find more resources in an agriculture-forest mosaic landscape than in a single land-cover dominated landscape (Vickery & Arlettaz 2012). In the same study site, Taiwan, Chen et al. (2022) also showed that landscape heterogeneity was relatively important in supporting high bird species richness in agricultural landscapes.

We also found strongly positive associations between Shannon’s diversity index and richness of all breeding birds, forest birds, and farmland birds at the 1-km scale (Fig. 2a4, 2b4, & 2c4), further supporting the important role of compositional landscape heterogeneity in driving species richness (Benton et al. 2003; Fahrig et al. 2011; Barbaro et al. 2021). On the other hand, configurational landscape heterogeneity, measured by edge density, showed no or even negative association with native species richness. These results imply that care is needed in how compositionally heterogeneous landscapes are designed in Taiwan, because fragmented landscapes that also have high edge density may lead to neutral or even negative conservation outcomes. Also, while increasing compositional heterogeneity might be a useful overall guiding principle, it must be borne in mind that particular species of very high conservation concern, or those with specialist requirements are likely to need bespoke conservation strategies (Pickett & Siriwardena 2011; Katayama et al. 2014).

### 4.2 Habitat amount is important, but the effect varies among scales and taxa

We found strongly positive associations between habitat amount and richness of forest and farmland birds, but the patterns differed among spatial scales and species groups. For forest birds, the proportion of forest cover was strongly positively associated with richness at both local and landscape scales (Fig. 2b1, 2b5), but there was a neutral effect of edge density (Fig. 2b3). Conforming with predictions from the habitat amount hypothesis (Fahrig 2013), the influence of habitat amount was more important than configurational heterogeneity for forest birds. From finer to broader scales, canopy of forest stands and wide forest patches both benefit forest birds.

Also in Taiwan, Lin et al. (2019) found relatively high bird species richness present in both native forest and coniferous plantation patches in a 50-hectare agriculture-forest mosaic landscape, perhaps because of the similar vegetation structure between them. Huang and Ding (2017) reported that remnant trees that provide abundant food attract avian frugivores from surrounding vegetation into restored forest sites. These results suggest that not only continuous large forest patches are important, but that high forest cover at fine scales can also support high species richness in Taiwan. In contrast, there was significant association between proportion of farmland cover and richness of farmland birds at the 100-m scale, but not at the 2-km scale. For farmland birds, the positive effect of habitat amount was only present at the local scale, while at the landscape scale, the negative effect of configurational heterogeneity (Fig. 2c3) was stronger than that of habitat amount (Fig. 2c6). Our results reflect that the relationship between landscape heterogeneity and richness of birds is scale-dependent and differs among species groups (Vickery & Arlettaz 2012). On landscape management, our results suggest that agricultural landscape management actions for farmland birds are perhaps best focused on a site-based scale.

### 4.3 Management strategies for introduced birds

Edge density was the only variable positively associated with the richness of introduced birds (Fig. 5-2d3), suggesting that introduced birds in Taiwan prefer a fragmented landscape. Indeed, myna and starling species including Javan Myna *Acridotheres javanicus*, Common Myna *Acridotheres tristis*, and Black-collared Starling *Gracupica nigricollis* are the main introduced species in Taiwan (Ding et al. 2020) and prefer human dominated fragmented landscapes, such as built-up areas and farmlands (Wang 2014). Introduced birds appear to avoid high coverage of forests at both local and landscape scales (Fig. 5-2d1, 5-2d5). Maintaining landscapes with intact, non-fragmented forest patches even at local scales might help to restrict the further expansion of introduced species.

### 4.4 Taiwan’s agri-environmental scheme

The results of this study could support the evidence-based development of agri-environmental schemes in Taiwan, suggesting that key actions are to improve compositional heterogeneity while minimising habitat fragmentation, and increase habitat amount at appropriate scales depending on the target species group in particular locations. Agri-environment schemes aim to decrease the negative impacts of agricultural management to improve the environmental quality of farmlands, including protecting natural habitats, preventing population decrease and improving ecosystem services (Kleijn & Sutherland 2003), but the details vary among countries. In Europe, it often focusses on managing the landscapes surrounding farmlands (Batáry et al. 2015). The US government provides conservation funding to landowners for improving conservation on farmlands (Lambert et al. 2007). Moreover, in Japan, the “Satoyama Initiative” aims to manage landscapes to harmonise both production and conservation (Katoh et al. 2009; Morimoto 2011). Unlike many other cases around the world, Taiwan is a mountainous island with limited flat land and intensive land use competition among sectors (Chung 2019). Thus, appropriately managing farmland for biodiversity be a central strategy for mitigating land use competition between agricultural management and biodiversity conservation. This study had identified a series of relationships between heterogeneity of agricultural landscapes and local biodiversity that vary among spatial scale. Crucial next steps are to conduct the similar studies on other taxa, and for single species of particular conservation concern to further fine tune agricultural landscape management in Taiwan.

## Declarations

### Ethical Approval

not applicable

### Availability of data and materials

not applicable

## References

Andrén H. 1994. Effects of habitat fragmentation on birds and mammals in landscapes with different proportions of suitable habitat: a review. Oikos 71: 355—366. 10.1111/1365-2664.13885

Barbaro L et al. 2021. Organic management and landscape heterogeneity combine to sustain multifunctional bird communities in European vineyards. Journal of Applied Ecology, 58, 1261–1271. 10.1111/1365-2664.13885

Batáry P, Dicks LV, Kleijn D, Sutherland WJ. 2015. The role of agriDenvironment schemes in conservation and environmental management. Conservation Biology 29(4): 1006—1016.

Bates D, Mächler M, Bolker B, Walker S (2015). “Fitting Linear Mixed-Effects Models Using lme4.” Journal of Statistical Software, 67(1), 1–48. doi:10.18637/jss.v067.i01.

Benton TG et al. 2003. Farmland biodiversity: is habitat heterogeneity the key? Trends in Ecology & Evolution, 18(4), 182–188.

Burnham KP, Anderson DR (2002) Model selection and multimodel inference: a practical information-theoretic approach. Second edition. New York: Springer-Verlag.

Chang SE. 2013. Blue Magpie TEAgriculture: Eco-tea Cultivation and Participatory Farming in Pinglin Satoyama, Taiwan. Procedia - Social and Behavioral Sciences, 101, 14–22.

Chen S-H et al. 2022. An empirical and expert-knowledge hybrid approach to implement farmland habitat assessment for birds. Conservation Science and Practice, 4, e12760. 10.1111/csp2.12760

Chung L-N. 2019. Land use competition of illegal industrial factories on Taiwan’s farmlands. Land Issues Research Quarterly, 18(3), 118–129.

DeFries, R., Rovero, F., Wright, P., Ahumada, J., Andelman, S., Brandon, K.,… & Liu, J. 2010. From plot to landscape scale: linking tropical biodiversity measurements across spatial scales. Frontiers in Ecology and the Environment, 8(3), 153–160.

Ding, T.-S., C.-S. Juan, R.-S. Lin, Y.-J. Tsai, J.-L. Wu, J. Wu and Y.-H. Yang. 2020. The 2020 TWBF Checklist of the Birds of Taiwan. Taiwan Wild Bird Federation. Taipei, Taiwan.

Fahrig L, Nuttle WK. 2005. Population ecology in spatially heterogeneous environments. In: Ecosystem Function in Heterogeneous Landscapes (eds Lovett GM, Jones CG, Turner MG & Weathers KC). Springer-Verlag, New York, 95—118.

Fahrig L et al. 2011. Functional landscape heterogeneity and animal biodiversity in agricultural landscapes. Ecology Letter, 12, 101–112.

Fahrig L. 2013. Rethinking patch size and isolation effects: the habitat amount hypothesis. Journal of Biogeography, 40, 1649—1663.

Fischer J, Lindenmayer DB. 2007. Landscape modification and habitat fragmentation: a synthesis. Global Ecology and Biogeography, 16, 265—280.

Foley JA, Ramankutty N, Brauman KA, Cassidy ES, Gerber JS, Johnston M, Mueller ND, O’Connell C, Ray DK, West PC, Balzer C, Bennett EM, Carpenter SR, Hill J, Monfreda C, Palasky S, Rockstörm J, Sheehan J, Siebert S, Tilman D, Zaks DPM. 2011. Solutions for a cultivated planet. Nature 478: 337–342.

Fox J, Weisberg S. 2019. An R Companion to Applied Regression, Third edition. Sage, Thousand Oaks CA. https://socialsciences.mcmaster.ca/jfox/Books/Companion/.

Gabriel D et al. 2010. Scale matters: the impact of organic farming on biodiversity at different spatial scales. Ecology Letters, 13, 858–869. 10.1111/j.1461-0248.2010.01481.x

Guerrero-Pineda C et al. 2022. An investment strategy to address biodiversity loss from agricultural expansion. Nature Sustainability, 5, 610–618. 10.1038/s41893-022-00871-2

Haddad NM, Gonzalez A, Brudvig LA, Burt MA, Levey DJ, Damschen EI. 2017. Experimental evidence does not support the Habitat Amount Hypothesis. Ecography 40: 48–55

Huang, J.-Y and T.-S. Ding. 2017. Remnant trees and surrounding vegetation determine avian frugivore visitation of restored forest sites in Taiwan. Forest Ecology and Management, 294, 20–26. 10.1016/j.foreco.2017.03.023

Katayama N, Amano T, Naoe S, Yamakita T, Komatsu I, et al. (2014) Landscape Heterogeneity–Biodiversity Relationship: Effect of Range Size. PLOS ONE 9(3): e93359. 10.1371/journal.pone.0093359

Katoh K, Sakai S, Takahashi T. 2009. Factors maintaining species diversity in satoyama, a traditional agricultural landscape of Japan. Biological Conservation 142: 1930—1936.

Kirk DA, Lindsay KF, Brook RW. 2011. Risk of agricultural practices and habitat changes to farmland birds. Avian Conservation and Ecology, 6(1): 5. 10.5751/ACE-00446-060105

Kleijn D, Sutherland WJ. 2003. How Effective are European Agri-Environment Schemes in Conserving and Promoting Biodiversity? Journal of Applied Ecology 40(6):947–969.

Ko, JC-J et al. 2017. Point count sampling data from the Taiwan Breeding Bird Survey. Taiwan Journal of Biodiversity, 19(4), 243–254.

Kremen C & Merenlender AM. 2018. Landscapes that work for biodiversity and people. Science, 362(6412), eaau6020.

Lambert DM, Sullivan P, Claassen R, Foreman L. 2007. Profiles of US farm households adopting conservation-compatible practices. Land Use Policy 24:72–88.

Lee K-C et al. 2016. Tailoring the Satoyama Initiative concepts to the national and local context: a case study of the collaborative planning process of a rice paddy cultural landscape in an indigenous community, Taiwan. In UNU-IAS and IGES (eds.) 2016, Mainstreaming concepts and approaches of socio-ecological production landscapes and seascapes into policy and decision-making (Satoyama Initiative Thematic Review vol. 2), United Nations University Institute for the Advanced Study of Sustainability, Tokyo.

Li C-F et al. 2013. Classification of Taiwan forest vegetation. Applied Vegetation Science 16, 698–719.

Lin, D.-L., S.-W. Fu, H.-W. Yuan, T.-S. Ding. 2019. Bird species richness in relation to land-use patch structure and vegetation structure in a forest-agriculture mosaic. Ornithological Science 18(2): 135–147.

Lin, D-L, Maron M, Amano T, Chang A-Y, Fuller RA. 2023. Using empirical data analysis and expert elicitation to identify habitat associations: The case of farmland-adapted birds in Taiwan. Ibis 165(3): 974–985.

Maxwell SL et al. 2016. Biodiversity: The ravages of guns, nets and bulldozers. Nature, 536, 143–145.

Mendenhall CD et al. 2014. Predicting biodiversity change and averting collapse in agricultural landscapes. Nature, 509, 213—217.

Merckx T, de Miranda MD, Pereira HM. 2019. Habitat amount, not patch size and isolation, drives species richness of macro-moth communities in countryside landscapes. Journal of Biogeography 46: 956–967.

Ministry of the Interior of Taiwan. 2015. Land use investigation of Taiwan. Online resources https://www.nlsc.gov.tw/LUI/FileDL/FileDLShow.aspx

Morelli F et al. 2013. Landscape heterogeneity metrics as indicators of bird diversity: Determining the optimal spatial scales in different landscapes. Ecological Indicators, 34, 372–379. 10.1016/j.ecolind.2013.05.021

Morimoto Y. 2011. What is Satoyama? Points for discussion on its future direction. Landscape and Ecological Engineering 7: 163—171.

Neumann JL et al. 2016. The compositional and configurational heterogeneity of matrix habitats shape woodland carabid communities in wooded-agricultural landscapes. Landscape Ecology, 31, 301–315.

Newton I. 2004. The recent declines of farmland bird populations in Britain: an appraisal of causal factors and conservation actions. Ibis, 146: 579–600. 10.1111/j.1474-919X.2004.00375.x

Perović D et al. 2015. Configurational landscape heterogeneity shapes functional community composition of grassland butterflies. Journal of Applied Ecology, 52, 505–513.

Power AG. 2017. Ecosystem services and agriculture: trade-offs and synergies. Philosophical Transactions of the Royal Society B 365: 2959—2971. 10.1098/rstb.2010.0143

R Core Team. (2020). R: A Language and Environment for Statistical Computing. R Foundation for Statistical Computing, Vienna, Austria. URL https://www.R-project.org/.

Riva F, Fahrig L. 2022. The disproportionately high value of small patches for biodiversity conservation. Conservation Letter, e12881.

Saura S. 2021. The Habitat Amount Hypothesis implies negative effects of habitat fragmentation on species richness. Journal of Biogeography, 48, 11–22. 10.1111/jbi.13958

Schindler S, Von Wehrden H, Poirazidis K, Wrbka T, Kati V (2013) Multiscale performance of landscape metrics as indicators of species richness of plants, insects and vertebrates. Ecological Indicators, 31, 41–48.

Stanton RL, Morrissey CA, Clark RG. 2018. Analysis of trends and agricultural drivers of farmland bird declines in North America: A review. Agriculture, Ecosystems and Environment 254: 244—254.

Su, H-J. 1992. A geographical data organization system for the botanical inventory of Taiwan. Botany Institute, Academia Sinica Monograph Series 12:23–36.

Tilman D, Fargione J, Wolff B, D’Antonio C, Dobson A, Howarth R, Schindler D, Schlesinger WH, Simberloff D, Swackhamer D. 2001. Forecasting agriculturally driven global environmental change. Science 292: 281–284.

Tscharntke T, Klein AM, Kruess A, Steffan-Dewenter I, Thies C. 2005. Landscape perspectives on agricultural intensification and biodiversity - ecosystem service management. Ecology Letters 8: 857–874.

Tulloch AIT, Barnes MD, Ringma J, Fuller RA, Watson JEM. 2016. Understanding the importance of small patches of habitat for conservation. Journal of Applied Ecology 53(2): 418—429.

Wang L-T. 2014. A study of the spatial-temporal dynamics and habitat models of native and invasive mynas in Taiwan. Master Thesis. Chinese Culture University.

Vickery, J. & R. Arlettaz. The importance of habitat heterogeneity at multiple scales for birds in European agricultural landscapes. In Fuller, R.J. (ed). 2012. Birds and habitat: relationships in changing landscapes. Cambridge University press.

Yang J et al. 2021. Effects of compositional and configurational heterogeneity of the urban matrix on the species richness of woody plants in urban remnant forest patches. Landscape Ecology 37, 619–632. 10.17632/h54cxnrb6w.1.

Zhang W, Ricketts TH, Kremen C, Carney K, Swinton SM. 2007. Ecosystem services and dis-services to agriculture. Ecological Economics 64: 253—260.

