## Supporting information for "Agricultural landscapes with high compositional heterogeneity support both forest and farmland birds in Taiwan"

Table S1 The list of explanatory variables and candidate models

| Variable | Candidate model |
| --- | --- |
| Forest_P_ | Richness ~ Elev + Area + Farm_P_ (L, 100-m) |
|  | Richness ~ Elev + Area + Farm_P_ (L & Q, 100-m) |
| Farm_P_ | Richness ~ Elev + Area + Farm_P_ (L, 100-m) |
|  | Richness ~ Elev + Area + Farm_P_ (L & Q, 100-m) |
| H’ | Richness ~ Elev + Area + Sha (0.5-km) |
|  | Richness ~ Elev + Area + Sha (1-km) |
|  | Richness ~ Elev + Area + Sha (2-km) |
| Edge | Richness ~ Elev + Area + Edge (0.5-km) |
|  | Richness ~ Elev + Area + Edge (1-km) |
|  | Richness ~ Elev + Area + Edge (2-km) |
| Forest_T_ | Richness ~ Elev + Area + Forest_T_ (L, 0.5-km) |
|  | Richness ~ Elev + Area + Forest_T_ (L & Q, 0.5-km) |
|  | Richness ~ Elev + Area + Forest_T_ (L, 1-km) |
|  | Richness ~ Elev + Area + Forest_T_ (L & Q, 1-km) |
|  | Richness ~ Elev + Area + Forest_T_ (L, 2-km) |
|  | Richness ~ Elev + Area + Forest_T_ (L & Q, 2-km) |
| Farm_T_ | Richness ~ Elev + Area + Farm_P_ (L, 0.5-km) |
|  | Richness ~ Elev + Area + Farm_P_ (L & Q, 0.5-km) |
|  | Richness ~ Elev + Area + Farm_P_ (L, 1-km) |
|  | Richness ~ Elev + Area + Farm_P_ (L & Q, 1-km) |
|  | Richness ~ Elev + Area + Farm_P_ (L, 2-km) |
|  | Richness ~ Elev + Area + Farm_P_ (L & Q, 2-km) |

Abbreviations: Forest_P_: mean proportion of forest cover (buffer of survey point), Farm_P_ : proportion of farmland cover (buffer of survey point), Edge: edge density, H’: Shannon’s diversity index, Forest_T_: proportion of forest coverage (buffer of survey transect), Farm_T_ : proportion of farmland cover (buffer of survey transect); L: linear term, Q: quadratic term

Table S2 Akaike information criterion (AIC) of each candidate variable of all breeding birds across spatial scales. Bold type indicates the selected variables with the lowest AICs.

| Variable | Candidate model | AIC |
| --- | --- | --- |
| Forest_P_ | Richness ~ Elev + Area + Farm_P_ (L, 100-m) | 2654.9 |
|  | **Richness ~ Elev + Area + Farm_P_ (L & Q, 100-m)** | **2622.2** |
| Farm_P_ | **Richness ~ Elev + Area + Farm_P_ (L, 100-m)** | **2669.4** |
|  | Richness ~ Elev + Area + Farm_P_ (L & Q, 100-m) | 2677.4 |
| H’ | Richness ~ Elev + Area + Sha (0.5-km) | 2638.0 |
|  | **Richness ~ Elev + Area + Sha (1-km)** | **2635.4** |
|  | Richness ~ Elev + Area + Sha (2-km) | 2674.0 |
| Edge | Richness ~ Elev + Area + Edge (0.5-km) | 2661.6 |
|  | **Richness ~ Elev + Area + Edge (1-km)** | **2655.3** |
|  | Richness ~ Elev + Area + Edge (2-km) | 2657.3 |
| Forest_T_ | Richness ~ Elev + Area + Forest_T_ (L, 0.5-km) | 2656.2 |
|  | **Richness ~ Elev + Area + Forest_T_ (L & Q, 0.5-km)** | **2635.1** |
|  | Richness ~ Elev + Area + Forest_T_ (L, 1-km) | 2659.4 |
|  | Richness ~ Elev + Area + Forest_T_ (L & Q, 1-km) | 2647.9 |
|  | Richness ~ Elev + Area + Forest_T_ (L, 2-km) | 2657.2 |
|  | Richness ~ Elev + Area + Forest_T_ (L & Q, 2-km) | 2650.4 |
| Farm_T_ | Richness ~ Elev + Area + Farm_P_ (L, 0.5-km) | 2679.9 |
|  | Richness ~ Elev + Area + Farm_P_ (L & Q, 0.5-km) | 2671.1 |
|  | Richness ~ Elev + Area + Farm_P_ (L, 1-km) | 2677.6 |
|  | Richness ~ Elev + Area + Farm_P_ (L & Q, 1-km) | 2666.8 |
|  | Richness ~ Elev + Area + Farm_P_ (L, 2-km) | 2685.2 |
|  | **Richness ~ Elev + Area + Farm_P_ (L & Q, 2-km)** | **2652.7** |

Abbreviations: Forest_P_: mean proportion of forest cover (buffer of survey point), Farm_P_ : proportion of farmland cover (buffer of survey point), Edge: edge density, H’: Shannon’s diversity index, Forest_T_: proportion of forest coverage (buffer of survey transect), Farm_T_ : proportion of farmland cover (buffer of survey transect); L: linear term, Q: quadratic term

Table S3 Akaike information criterion (AIC) of each candidate variable of forest birds across spatial scales. Bold type indicates the selected variables with the lowest AICs.

| Variable | Candidate model | AIC |
| --- | --- | --- |
| Forest_P_ | **Richness ~ Elev + Area + Farm_P_ (L, 100-m)** | **2186.1** |
|  | Richness ~ Elev + Area + Farm_P_ (L & Q, 100-m) | 2254.4 |
| Farm_P_ | **Richness ~ Elev + Area + Farm_P_ (L, 100-m)** | **2191.6** |
|  | Richness ~ Elev + Area + Farm_P_ (L & Q, 100-m) | 2192.2 |
| H’ | Richness ~ Elev + Area + Sha (0.5-km) | 2140.3 |
|  | **Richness ~ Elev + Area + Sha (1-km)** | **2131.8** |
|  | Richness ~ Elev + Area + Sha (2-km) | 2175.4 |
| Edge | Richness ~ Elev + Area + Edge (0.5-km) | 2168.8 |
|  | Richness ~ Elev + Area + Edge (1-km) | 2157.4 |
|  | **Richness ~ Elev + Area + Edge (2-km)** | **2152.3** |
| Forest_T_ | Richness ~ Elev + Area + Forest_T_ (L, 0.5-km) | 2102.5 |
|  | Richness ~ Elev + Area + Forest_T_ (L & Q, 0.5-km) | 2099.0 |
|  | Richness ~ Elev + Area + Forest_T_ (L, 1-km) | 2098.2 |
|  | Richness ~ Elev + Area + Forest_T_ (L & Q, 1-km) | 2121.5 |
|  | **Richness ~ Elev + Area + Forest_T_ (L, 2-km)** | **2085.8** |
|  | Richness ~ Elev + Area + Forest_T_ (L & Q, 2-km) | 2138.7 |
| Farm_T_ | Richness ~ Elev + Area + Farm_P_ (L, 0.5-km) | 2176.9 |
|  | Richness ~ Elev + Area + Farm_P_ (L & Q, 0.5-km) | 2171.1 |
|  | Richness ~ Elev + Area + Farm_P_ (L, 1-km) | 2123.9 |
|  | **Richness ~ Elev + Area + Farm_P_ (L & Q, 1-km)** | **2109.3** |
|  | Richness ~ Elev + Area + Farm_P_ (L, 2-km) | 2186 |
|  | Richness ~ Elev + Area + Farm_P_ (Q, 2-km) | 2.137.2 |

Abbreviations: Forest_P_: mean proportion of forest cover (buffer of survey point), Farm_P_ : proportion of farmland cover (buffer of survey point), Edge: edge density, H’: Shannon’s diversity index, Forest_T_: proportion of forest coverage (buffer of survey transect), Farm_T_ : proportion of farmland cover (buffer of survey transect); L: linear term, Q: quadratic term

Table S4 Akaike information criterion (AIC) of each candidate variable of farmland birds across spatial scales. Bold type indicates the selected variables with the lowest AICs.

| Variable | Candidate model | AIC |
| --- | --- | --- |
| Forest_P_ | Richness ~ Elev + Area + Farm_P_ (L, 100-m) | 1890.2 |
|  | Richness ~ Elev + Area + Farm_P_ (L & Q, 100-m) | 1904.6 |
| Farm_P_ | Richness ~ Elev + Area + Farm_P_ (L, 100-m) | 1977.1 |
|  | Richness ~ Elev + Area + Farm_P_ (L & Q, 100-m) | 2032.7 |
| H’ | Richness ~ Elev + Area + Sha (0.5-km) | 1800.8 |
|  | **Richness ~ Elev + Area + Sha (1-km)** | **1783.2** |
|  | Richness ~ Elev + Area + Sha (2-km) | 1878.7 |
| Edge | Richness ~ Elev + Area + Edge (0.5-km) | 1870.9 |
|  | **Richness ~ Elev + Area + Edge (1-km)** | **1865.7** |
|  | Richness ~ Elev + Area + Edge (2-km) | 1865.8 |
| *Forest_T_* | Richness ~ Elev + Area + Forest_T_ (L, 0.5-km) | 1878.7 |
|  | **Richness ~ Elev + Area + Forest_T_ (L & Q, 0.5-km)** | **1801.3** |
|  | Richness ~ Elev + Area + Forest_T_ (L, 1-km) | 1885.3 |
|  | Richness ~ Elev + Area + Forest_T_ (L & Q, 1-km) | 1816.9 |
|  | Richness ~ Elev + Area + Forest_T_ (L, 2-km) | 1885.5 |
|  | Richness ~ Elev + Area + Forest_T_ (L & Q, 2-km) | 1818.1 |
| *Farm_T_* | Richness ~ Elev + Area + Farm_P_ (L, 0.5-km) | 1888.2 |
|  | Richness ~ Elev + Area + Farm_P_ (L & Q, 0.5-km) | 1871.5 |
|  | Richness ~ Elev + Area + Farm_P_ (L, 1-km) | 1876.5 |
|  | Richness ~ Elev + Area + Farm_P_ (L & Q, 1-km) | 1879.0 |
|  | Richness ~ Elev + Area + Farm_P_ (L, 2-km) | 1890.2 |
|  | **Richness ~ Elev + Area + Farm_P_ (L & Q, 2-km)** | **1871.6** |

Abbreviations: Forest_P_: mean proportion of forest cover (buffer of survey point), Farm_P_ : proportion of farmland cover (buffer of survey point), Edge: edge density, H’: Shannon’s diversity index, Forest_T_: proportion of forest coverage (buffer of survey transect), Farm_T_ : proportion of farmland cover (buffer of survey transect); L: linear term, Q: quadratic term

Table S5 Akaike information criterion (AIC) of each candidate variable of introduced birds across spatial scales. Bold type indicates the selected variables with the lowest AICs.

| Variable | Candidate model | AIC |
| --- | --- | --- |
| Forest_P_ | **Richness ~ Elev + Area + Farm_P_ (L, 100-m)** | **1110.6** |
|  | Richness ~ Elev + Area + Farm_P_ (L & Q, 100-m) | 1119.0 |
| Farm_P_ | **Richness ~ Elev + Area + Farm_P_ (L, 100-m)** | **1139.0** |
|  | Richness ~ Elev + Area + Farm_P_ (L & Q, 100-m) | 1139.1 |
| H’ | Richness ~ Elev + Area + Sha (0.5-km) | 1116.1 |
|  | Richness ~ Elev + Area + Sha (1-km) | 1117.2 |
|  | **Richness ~ Elev + Area + Sha (2-km)** | **1112.1** |
| Edge | Richness ~ Elev + Area + Edge (0.5-km) | 1113.6 |
|  | Richness ~ Elev + Area + Edge (1-km) | 1111.3 |
|  | **Richness ~ Elev + Area + Edge (2-km)** | **1109.6** |
| Forest_T_ | Richness ~ Elev + Area + Forest_T_ (L, 0.5-km) | 1116.9 |
|  | Richness ~ Elev + Area + Forest_T_ (L & Q, 0.5-km) | 1113.3 |
|  | Richness ~ Elev + Area + Forest_T_ (L, 1-km) | 1116.8 |
|  | Richness ~ Elev + Area + Forest_T_ (L & Q, 1-km) | 1113.5 |
|  | Richness ~ Elev + Area + Forest_T_ (L, 2-km) | 1116.9 |
|  | **Richness ~ Elev + Area + Forest_T_ (L & Q, 2-km)** | **1113.2** |
| Farm_T_ | Richness ~ Elev + Area + Farm_P_ (L, 0.5-km) | 1110.3 |
|  | Richness ~ Elev + Area + Farm_P_ (L & Q, 0.5-km) | 1110.9 |
|  | Richness ~ Elev + Area + Farm_P_ (L, 1-km) | 1115.5 |
|  | Richness ~ Elev + Area + Farm_P_ (L & Q, 1-km) | 1112.4 |
|  | **Richness ~ Elev + Area + Farm_P_ (L, 2-km)** | **1100.7** |
|  | Richness ~ Elev + Area + Farm_P_ (L & Q, 2-km) | 1107.8 |

Abbreviations: Forest_P_: mean proportion of forest cover (buffer of survey point), Farm_P_ : proportion of farmland cover (buffer of survey point), Edge: edge density, H’: Shannon’s diversity index, Forest_T_: proportion of forest coverage (buffer of survey transect), Farm_T_ : proportion of farmland cover (buffer of survey transect); L: linear term, Q: quadratic term

Table S6 Variance Inflation Factor (VIF) scores of explanatory variables in the best model of each species group

| Variable | VIFs of each species group | | | |
| --- | --- | --- | --- | --- |
|  | Total birds | Forest birds | Farmland birds | Introduced birds |
| Forest_P_ | 2.63 | 3.49 | 2.50 | 2.43 |
| Farm_P_ | 1.74 | 2.71 | 1.81 | 1.86 |
| Elev | 1.61 | 1.83 | 1.70 | 1.74 |
| Area | 1.06 | 1.04 | 1.06 | 1.06 |
| H’ | 2.69 | 1.60 | 2.62 | 3.46 |
| Edge | 1.55 | 3.10 | 1.98 | 3.55 |
| Forest_T_ | 3.76 | 3.80 | 2.63 | 1.25 |
| Farm_T_ | 1.59 | 2.78 | 1.62 | 2.61 |

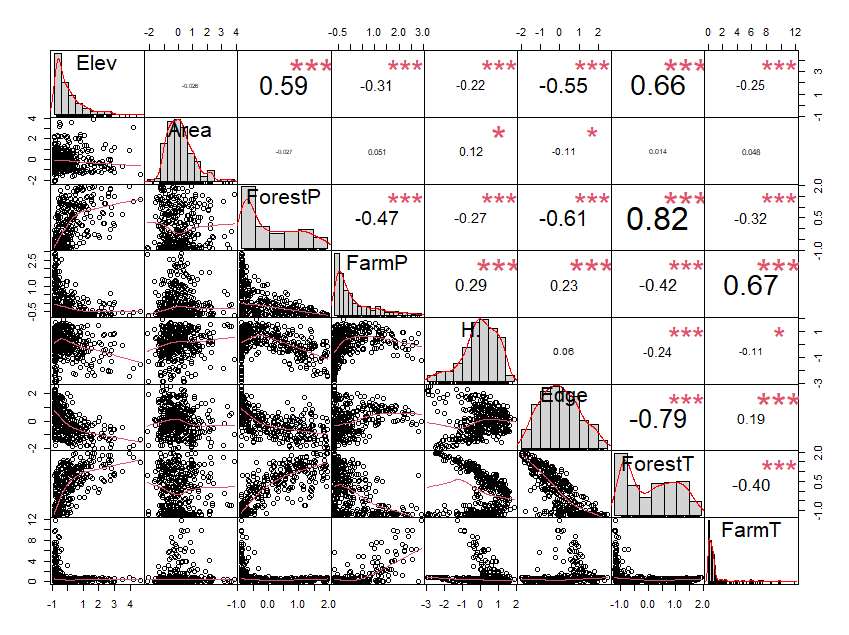

Fig. S1 The correlation matrix of the explanatory variables in the best model of total bird species richness.

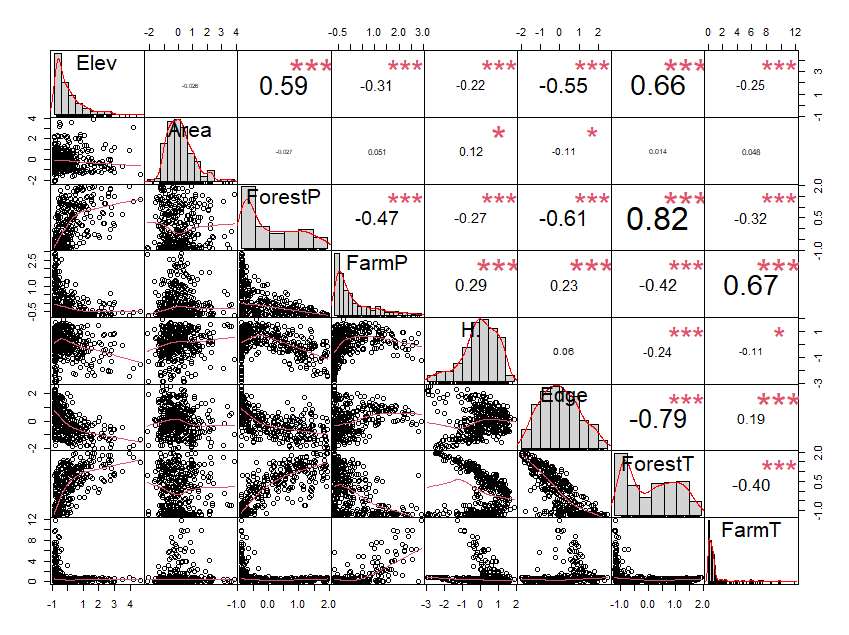

Fig. S2 The correlation matrix of the explanatory variables in the best model of forest bird species richness.

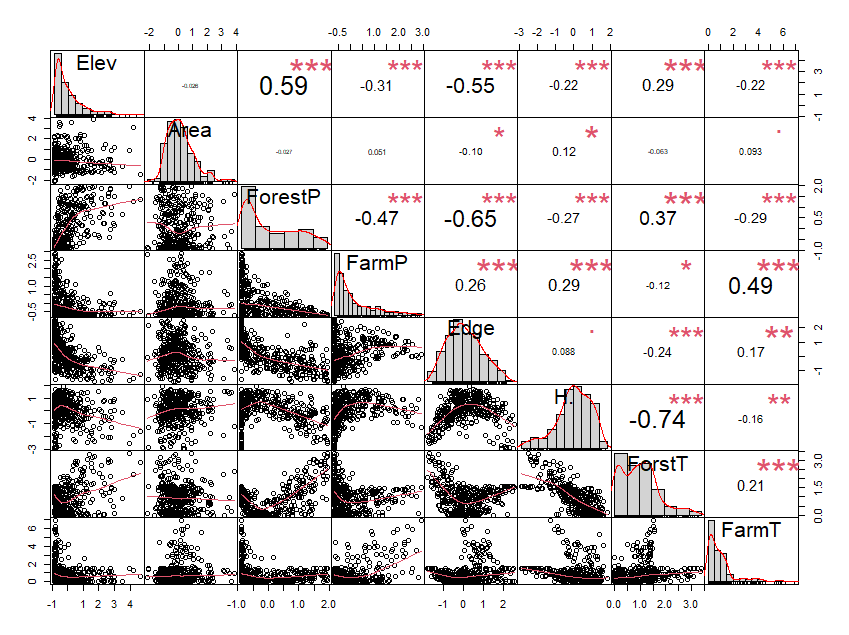

Fig. S3 The correlation matrix of the explanatory variables in the best model of forest bird species richness.

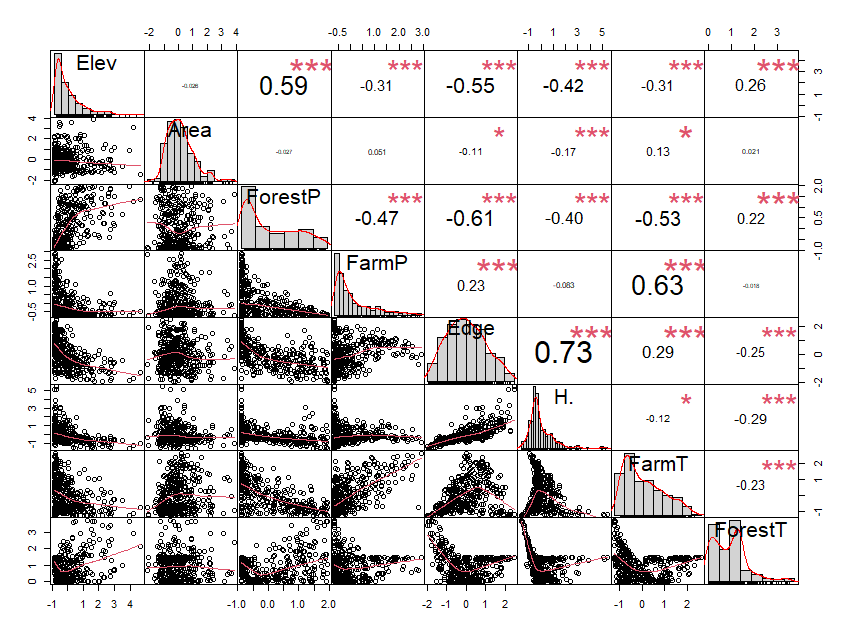

Fig. S4 The correlation matrix of the explanatory variables in the best model of forest bird species richness.
